# Adding 3Di characters to amino acid datasets can improve resolution, but the effect is weaker in shorter and alpha-helical proteins

**DOI:** 10.1101/2025.06.30.662300

**Authors:** Matthew S. Fullmer, Caroline Puente-Lelievre, Nicholas J. Matzke

## Abstract

The recent introduction of Foldseek’s 3Di character alphabet to encode 3D protein structure has opened up new possibilities for structural phylogenetics. These characters, like protein structure, are more conserved than amino acids, raising the possibility of better resolution of very deep branches on the tree of life. As 3Di characters have a 20-letter alphabet, they are readily treatable with off-the-shelf algorithms for model-based phylogenetic inference and related methods such as bootstrapping. However, it remains to be seen if 3Di phylogenies are broadly more resolved than sequence-based phylogenies. We present data using samples from nine protein superfamilies showing that 3Di combines with sequence to produce better resolved phylogenies than either sequence or 3Di alone. We also show that information-theoretic measures, applied to superfamily alignments, significantly correlate with resolution in phylogenies derived from these alignments. Further, we identify the proportion of alpha helices in proteins as a major driver in reducing the information carried by 3Di character alignments, explaining the relatively poor performance of 3Di characters on superfamilies with highly-conserved structure but high alpha helical content. Our results provide encouragement for the further use of 3Di to address challenging questions in deep history, but also sound a note of caution about which proteins it is most suitable for.

**SIGNIFICANCE:** 3Di characters have been suggested as a method to generate well-resolved deep phylogenies. Our results show that 3Di characters combined with sequences can improve resolution in the deepest nodes of protein superfamily trees. However, our results also show that 3Di characters may not be suitable for all protein types.

## INTRODUCTION

It has long been understood that protein shape may contain phylogenetic information [1]. Protein 3D structure is typically more conserved than amino acid (AA) sequences [2,3]. Attempts have been made to use structural distance measures for neighbor-joining phylogenetic inference (e.g. Lundin et al., 2012 [4]; see Puente-Lelievre et al. 2024a [5] for a review), however these approaches lack an evolutionary model and the associated advantages of probabilistic approaches, such as uncertainty measures. An extension of the structural distances approach [6] uses simulated conformational variability to create a pseudo-support tree set, however it is computationally very expensive. An alternative approach is to encode protein structure as a series of discrete characters, an approach which became possible with the protein structure alphabet developed as part of the Foldseek program [7]. The alphabet is termed “3Di” as it encodes highly conserved 3D tertiary interactions in a protein. The Foldseek program translates any input protein structure model into a series of 3Di characters, one for each residue in the protein backbone. The 3Di alphabet conveniently has 20 characters, enabling usage of BLAST and other standard bioinformatics algorithms for homology search. As the 3Di characters are highly conserved, Foldseek is able to conduct high-sensitivity deep homology searches on databases of structure models in and beyond the “Twilight Zone” of highly decayed sequence similarity. This capability is particularly powerful with the availability of databases of AlphaFold-derived structural predictions.

Beyond deep homology search, having a discrete 3Di structural alphabet that is more conserved than AAs opens new possibilities for structural phylogenetics [5,8–10]. We focus on use of 3Di characters to enhance model-based phylogenetic inference, making use of highly-conserved protein structure data while allowing all the advantages of probabilistic methods, including statistical model comparison and standard measures of statistical confidence such as bootstrapping. While early reports are promising [5,11], the suitability of 3Di characters for phylogenetics needs further exploration.

One of the attractions of structural phylogenetics is the prospect of better resolving the earliest branches of protein evolution. These early branches represent some of the earliest evolution in life’s history. In many protein superfamilies, they would predate the Last Universal Common Ancestor (LUCA). We investigate whether 3Di characters are likely to help with such problems by comparing various measures of phylogenetic resolution on datasets of ancient protein families that have diverged into the Twilight Zone, when making use of AA, 3Di, or AA+3Di character data.

To begin to explore this question, we estimated alignments and phylogenies for 9 protein superfamilies, comparing AA-only, 3Di-only, or AA+3Di datasets with several measures of statistical resolution. We also looked for differences in resolution between deep and shallow branches. We observe that, across the 9 protein superfamilies, there is a substantial and statistically significant increase in Tree Certainty and related measures of phylogenetic resolution for AA+3Di datasets over AA-only or 3Di-only datasets. This suggests that 3Di characters are a useful addition in the Twilight Zone, even though not all deep divergences are resolved. However, the results vary by superfamily, with highly alpha-helical protein superfamilies not showing substantial resolution improvements with the addition of 3Di characters.

We expanded our analysis to examine possible correlates that might be responsible for the variation in resolution between superfamilies. Our results suggest that proteins containing large proportions of alpha-helices may result in drops in the information content of the 3Di-coded alphabet, and a commensurate drop in phylogenetic resolution.

## RESULTS

### Comparison of AA-only, 3Di-only, and AA+3Di analyses across 9 superfamilies

Overall, combining AA+3Di offers significant improvements in tree resolution (**Figure 1**). The AA+3Di partitioned dataset gained ∼11% in average bootstrap percentage over AA-only datasets, and ∼0.17 TC/TCA. All of these gains were statistically significant (bootstrap p = 0.019, TC/TCA p = 0.00053 and p = 0.00040). Also encouraging, the improvements were not observed only in shallow nodes. For mean bootstrap percentages on deeper nodes (**Figure 2**), AA+3Di gained 22% over AA-alone (P=0.078, not significant) and 36% over 3Di alone (P=0.027). For mean bootstrap percentages on younger nodes, AA+3Di gained 18% over AA-alone (P=0.027, significant) and 28% over 3Di alone (P=0.0047, significant). While the improvement seems more consistent on younger nodes, overall, combining AA+3Di appears to be improving resolution on average. For superfamily descriptions and summary statistics, see **Table S1**.

**Figure 1.**
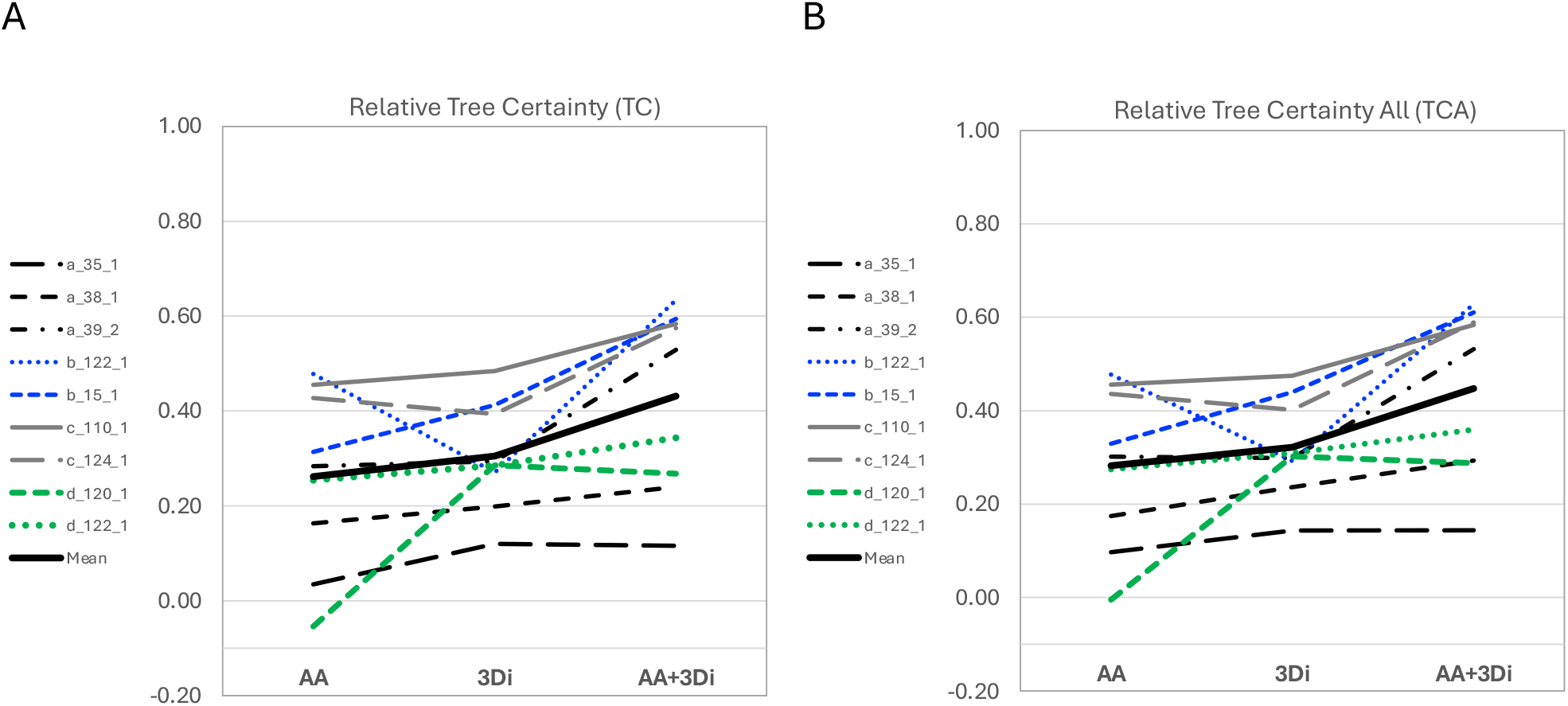
Comparison of full phylogenies of AA-only, 3Di-only, and AA+3Di analyses across all 9 superfamilies. A) Relative Tree Certainty; B) Relative Tree Certainty All.

**Figure 2.**
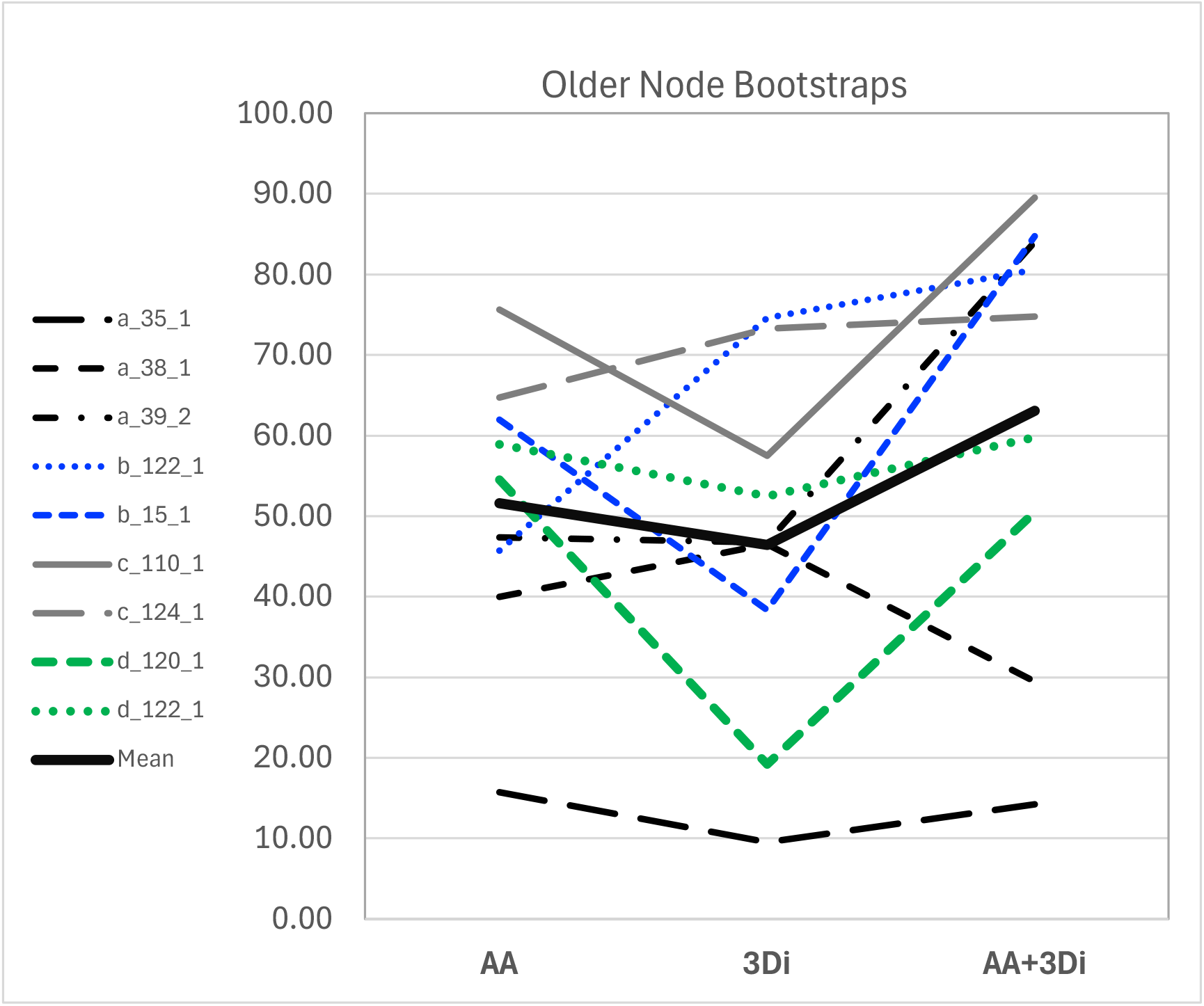
Comparison of only deeper nodes of AA-only, 3Di-only, and AA+3Di analyses across all 9 superfamilies. Measure of resolution is average bootstrap value per node.

These average improvements do not apply to all datasets, as can be observed in Figures 1-2, where several datasets show marginal or no improvement from adding 3Di characters. In particular, two of the alpha-helical superfamilies, a.35.1 (lambda repressor-like DNA-binding domains) and a.38.1 (HLH, helix-loop-helix DNA-binding domain) show the lowest overall resolution in TC and TCA terms, and little or no improvement in AA+3DI analyses. For a.38.1, AA+3Di analysis actually shows a decline in mean bootstrap percentage for older nodes. In order to explore the reasons why, we examined correlations of resolution measures with structural features of the superfamilies.

### Shorter and more alpha-helix-heavy proteins have reduced phylogenetic resolution

The two poorly-performing alpha-helical superfamilies, a.35.1 and a.38.1, had mean lengths of 85 and 73 AAs, respectively. The other superfamily that showed no improvement in older node bootstrap resolution (d.120.1, Figure 2) was also the third shortest in average length (88 AAs). Average sequence length was significantly positively correlated with our measures of phylogenetic resolution among the 9 superfamilies (**Figure 3**). Having more sequence length simply tends to carry more information [12].

**Figure 3.**
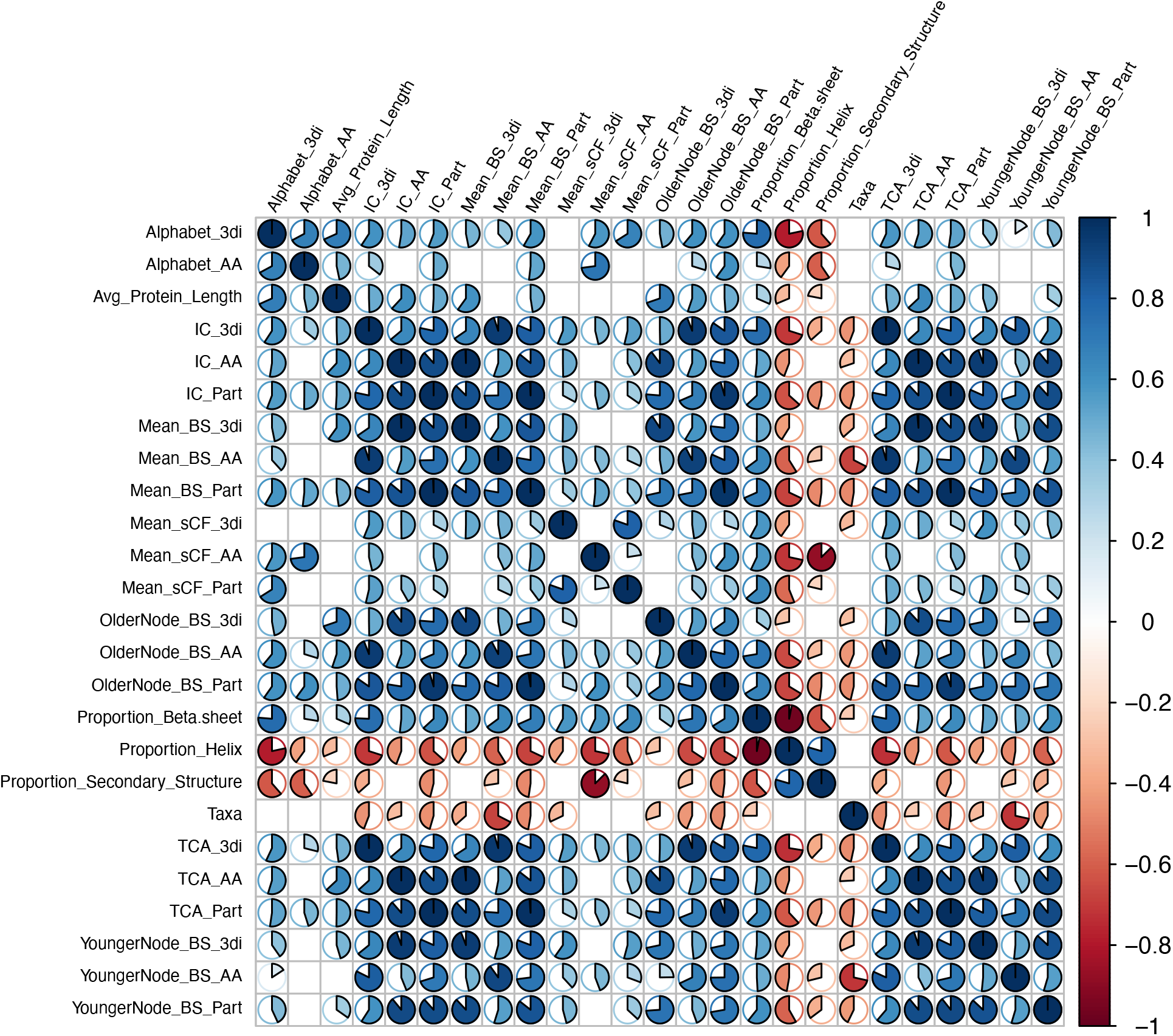
Pearson correlations of protein superfamily characteristics and phylogeny support values. Display cutoff is p = 0.01. Smaller alphabet size is significantly correlated with reduced resolution. Similarly, the proportion of the protein composed of helices is correlated with smaller alphabets and reduced resolution.

The a.35.1 and a.38.1 superfamilies are also heavily alpha-helical. They are composed of an average of 62% and 74% percent alpha helices, respectively (Table 1). As a result, the 3Di alignments are dominated by only a few character states common in alpha-helices, with V being most dominant and substantially larger than D, the second-most frequent character state (Table S2). For superfamily a.35.1, the frequency of V=33.25%, and D=13.64%. For a.38.1, V=62.42%, with D=11.74%. For the better-resolved a.39.2, V=29.08% and D=13.81%, but this still higher than the average across all superfamilies (mean V=25.1%), or the next-most V-enriched superfamily, the relatively poorly-resolved d.120.1 (V=25.05%, with D=14.87% coming second). For the alpha-helical superfamilies, the leading three states comprise 56.64%, 82.97%, and 56.33% of all characters, respectively. In contrast, for all the other superfamilies, the highest total frequency for the leading three states is 46.66% (d.120.1).

**Table 1.** Summary statistics on secondary structure proportions and effective alphabet sizes for the 9 studied superfamilies.

| Superfamily | Taxa Meeting Criteria | Mean Protein Length | Proportion Alpha Helix | Proportion Beta Sheet | Proportion Secondary Structure | Effective Alphabet Size (AAs) | Effective Alphabet Size (3di) |
| --- | --- | --- | --- | --- | --- | --- | --- |
| a.35.1 | 32 | 85.0 | 0.623 | 0.025 | 0.648 | 16.90 | 9.12 |
| a.38.1 | 7 | 73.0 | 0.744 | 0.000 | 0.744 | 15.14 | 3.86 |
| a.39.2 | 13 | 126.5 | 0.622 | 0.015 | 0.637 | 18.16 | 8.77 |
| b.122.1 | 13 | 136.5 | 0.245 | 0.313 | 0.557 | 17.75 | 15.57 |
| b.15.1 | 11 | 109.3 | 0.094 | 0.433 | 0.527 | 16.82 | 11.85 |
| c.110.1 | 8 | 144.4 | 0.258 | 0.397 | 0.655 | 17.43 | 15.65 |
| c.124.1 | 23 | 241.5 | 0.387 | 0.201 | 0.588 | 17.62 | 15.07 |
| d.120.1 | 15 | 87.9 | 0.340 | 0.211 | 0.550 | 17.55 | 12.52 |
| d.122.1 | 26 | 191.1 | 0.355 | 0.273 | 0.627 | 17.24 | 15.38 |

### Smaller alphabet sizes significantly correlate with lower support values

The combination of short and helix-heavy proteins are information-poor alignments - particularly notable in the 3Di data. Calculating entropies to estimate the effective alphabet sizes of the empirical character frequencies reveals that the 3Di sequences, despite having 20 character states, are functionally using an alphabet of only size 9.1 and 3.9 for a.35.1 and a.38.1. The non-alpha-helical superfamilies have alphabet sizes ranging from 11.9-15.7 (Table 1; Figure 4). For amino acids, however, effective alphabet sizes remain between 15-18 across all superfamilies (Table 1). Pearson correlations (**Figure 3**) and linear regressions (**Figure 4**) found significant relationships between proportion of alpha helix, 3Di alphabet size, and the resolution of 3Di-derived phylogenies across the superfamilies.

**Figure 4.**
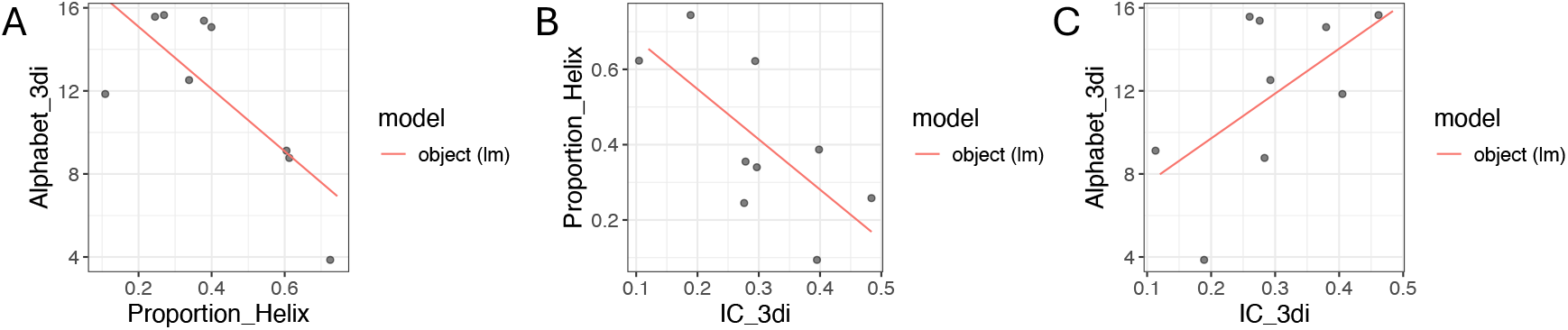
Linear Regressions of relationships between phylogeny resolution (IC (TC) 3Di), 3Di Alphabet size, and Proportion of Helices. A) 3Di Alphabet vs Proportion of Helix. B) 3Di TC values vs Proportion of Helices; C) 3Di IC (TC) values vs 3Di Alphabet sizes. Increasing proportion of helices appears to drive down the 3di alphabet size. In turn, smaller 3Di alphabets appear to drive down phylogenetic resolution.

### Helix proportion is significantly anti-correlated with both alphabet size and support values

Helix proportion was by far the largest anti-correlation with effective alphabet size. This was far more emphatic for the 3Di alphabets than AA (although the link is strong in both). This aligns with the conclusion that helix proportion is a driver of reduced alphabets (**Figure 4**).

We were concerned that this result could be a bias from including three alpha-only superfamilies and only 2 of each other fold type. We tested this by removing one, two, or all of the alpha-only superfamilies and calculating linear regressions between helix proportion and alphabet size. While the effect size dropped, including only two still shows a significant relationship, suggesting that this is a real factor (**Table 2**).

**Table 2.**
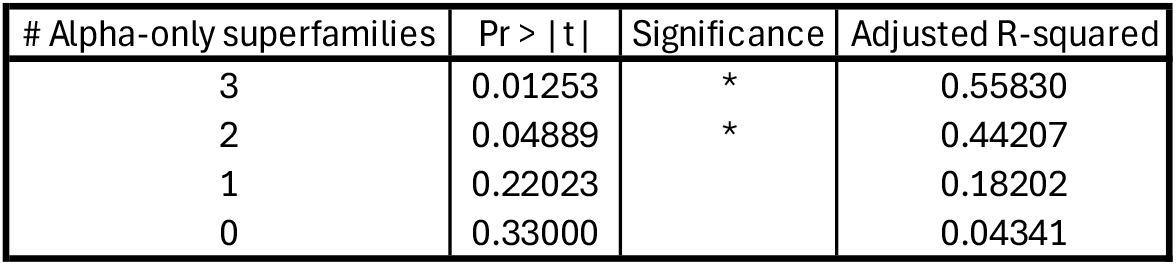
Linear regressions of helix proportion vs 3Di alphabet size. Rows vary the number of alpha-only superfamilies in the regression. When the number of alpha-only superfamilies is reduced to 2, the same number of superfamilies in the three other fold-types, the relationship remains strong and significant.

Conversely, beta-sheet proportion is positively correlated with alphabet size, as well as higher average support values. As both alphabet sizes are inversely correlated with support values, it seems reasonable to suggest that enrichment in helices, and perhaps paucity of beta-sheets, is a major driver behind poorer resolution in alpha-rich superfamilies.

## DISCUSSION

We present evidence that 3Di combined with sequence tends to improve overall resolution, as well as resolution in deeper nodes when compared to sequence or 3Di alone. This bolsters the promise of using 3Di in model-based phylogenetics [5,13], although it does not suggest that 3Di will resolve every ancient relationship.

We also present evidence that superfamilies with shorter protein lengths and enrichment in alpha-helices are less suitable for 3Di-based phylogenetics. These two factors combine to reduce the effective alphabet size of their alignments, thereby reducing the amount of information conveyed. These reductions in information content significantly correlate with reduced phylogenetic resolution. These results offer a note of caution, suggesting that researchers doing deep phylogenies should calibrate their expectations by considering the characteristics of proteins they are studying before deploying 3Di characters.

Delving more deeply into the results on poorly-performing alpha-helix-dominated proteins, we suggest that the main limitations on 3Di performance are poor character state diversity and domination by a few states: V and D, and to a lesser extent, L, S and P). Further investigation in other superfamilies is important to further test the degree to which this is a generalisable result; however, anecdotally, we have observed many 3Di alignments where the alpha-helical regions are dominated by these states).

In this work, we focused on the resolution of inferred phylogenies. We did not focus on the actual topologies the methods produced, as we do not know the “true” history of any of these ancient superfamilies. “Ground truthing” a new set of phylogenetic characters against an already known phylogeny is possible in simulations and in certain laboratory experiments, but these are difficult options to enact as a valid test would have to include realistic protein structure evolution over deep time, as well as sequence divergence into the Twilight Zone. However, these problems not unique, as similar problems are present for many sources of phylogenetic characters (fossil morphology, developmental characters, behaviour, etc.). Nevertheless, even in the absence of ground-truthing, being able to deploy methods from the suite of model-based statistical phylogenetics, such methods to measure uncertainty (bootstrapping, posterior probabilities, etc.), constitutes a major advantage, enabling statistical studies that allow us to at least quantify the precision that structural phylogenetics may achieve, and compare that to more traditional methods.

## METHODS

### SCOP superfamilies

See **Table S1** for full listing of the superfamilies used in this analysis. SCOP 1.75 supplied the categorization information [14,15]. Superfamilies were initially selected with the goal of evenly sampling the 4 major classes containing alpha helix and beta sheet-containing proteins (alpha-only, beta-only, a+b, and a/b). Another goal was to sample across a variety of size ranges, such that we could interrogate the correlation of sequence length and secondary structure composition with phylogenetic resolution. PDB files for each selected superfamily were downloaded from the PDB database. Each prospective superfamily was processed in a manner based on the filtering and clustering described for STRUCTOME [16]. All proteins below 50 AAs and above 350 AAs were excluded. UCLUST was used to prune the dataset, using centroid clustering at 90% identity to identify a single sequence to represent a group of similar sequences [17], making phylogenetic inference for each superfamily more computationally tractable. However, clustering also meant that there was no guarantee that each family in the superfamily would have a representative, or that component families would have equal or proportional representation. From the superfamilies that, after processing, retained at least 4 remaining members, 9 were chosen: three “a,” and two each of “b/c/d.”

### Secondary structure annotation

DSSP, as implemented in XSSP [18,19] was used to quantify the secondary structure proportions of each sampled sequence. All three helix types (H, G, I) were combined into a single proportion. Likewise, both beta-sheet types (B, E) were also combined into a single proportion. The metric “total secondary structure proportion” represents the sum of helix and sheet proportions.

### Tree Certainty Measures

Uncertainty of tree collections (e.g. bootstrap support sets) was measured using Internode Certainty (IC). IC represents a quantification of the level of disagreement in a support set for a particular node in a phylogeny; IC decreases as the bootstrap value drops and as the frequency of the most frequent dissenting topology increases. Tree Certainty (TC) is the average Internode Certainty score across all internal branches [20] (see a brief summary of the terminology and metrics in **Supplemental Material**). These scores were calculated by mapping support tree sets against reference trees, as implemented in RAxML v8.1 [21]. The related Internode Certainty All (ICA) score was also calculated. The ICA considers not only the most frequent dissenting topology at a branch, but also whether dissenting trees agree or disagree with each other. I.e., a bipartition supported by 51% of bootstrap trees would score much higher if the conflicting samples represented 49 different alternatives than it would if there were only a single alternative. The Tree Certainty All (TCA) score is the average of ICA values across internal branches, representing an assessment of overall conflict in the support set.

### Mean bootstrap values

Bootstrap values across the branches of each tree were extracted using custom R [22] scripts, using the R library *ape* [23]. Scripts and data are available in links supplied in Supplemental Material.

### Amino acid (AA) phylogenetic inference

For the analyses presented here, Foldmason [9] was employed to create structurally-informed AA superfamily alignments. Maximum likelihood phylogenetic interference was performed in IQ-TREE2, with substitution models selected by IQ-TREE’s model finder using the BIC criterion [24].

### 3Di phylogenetic inference

Foldmason was employed to create structurally-informed joint 3Di and AA alignments [9]. Maximum likelihood interference of phylogenies was done in IQ-TREE2 using the Foldseek 3Di rate matrix as in in Puente-Lelievre et al. (2024a), and using IQ-TREE’s model finder to choose base-frequency and among-site rate variation models [24].

### Partitioned analysis of AA+3Di datasets

Partitioned analyses were performed with IQ-TREE2 with the “-p” option [24]. The models for each partition were determined as above for the AA-only and 3Di-only datasets.

### 3Di rate matrix

The rate matrix from [5] was used with all 3Di alignments. Four superfamilies (a.35.1, a.38.1, a.39.2, and b.122.1) were tested using the alternate rate matrices presented in [11]. Their average bootstrap percentages were both lower (51% vs 45%, and 49%). Linear regression found the difference to be weakly significant (p = 0.018 and 0.0299). This indicated resolution would be better with the Foldseek-derived rate matrix (**Table S3**).

### Effective Alphabet Size

Raw character frequencies from alignments were obtained from IQ-TREE2 [24]. Shannon entropies were calculated for each alignment, summed and converted into an effective alphabet size [25].

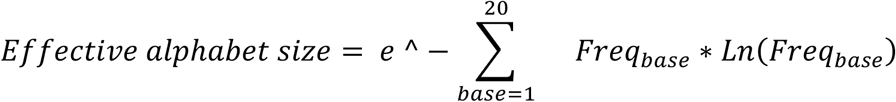

### sCF

Site concordance factors were calculated in IQ-TREE2 with the -blfix option [24,26]. For3Di alignments, the model was fixed to the3Di rate matrix. IQ-TREE was allowed to re-estimate AA models.

### Correlogram

The R function *cor* was used to calculate Pearson correlations. *cor*.*test* was used to calculate significance. Linear models were calculated with R’s *lm* function. The R library *corrplot* [27] visualized the correlogram. Only correlations with significance p =< 0.01 were displayed.

### Old/Young nodes

In order to judge some nodes to be relatively older than others, a rooted tree is needed. While BEAST automatically estimates the root position, IQ-TREE does not. Therefore, for IQ-TREE phylogenies, we used midpoint rooting in R package *phytools* [28], which places the root in the middle of the longest path through the tree. Midpoint rooting is known to be uncertain and can be affected by artefacts such as long branches, but outgroups are typically not available for such ancient divergences, and at least midpoint roots plausibly place the root far from the tips of the tree, typically separating major named groups (e.g. protein families) into separate clades. For our purpose, which was merely to roughly distinguish statistical resolution at deeper versus shallower nodes, this was adequate. For each node in a tree, the root-to-node distance was calculated with R library *castor* [29], 2018). The median of these root-to-node distances was calculated for each tree, and nodes closer to root than median were considered “deep,” while those above the median were “young.” Mean support values are calculated as above.

## Acknowledgements

The work of MF was supported by 22-UOA-250. The work of NJM, CPL, and their lab group was primarily supported by HFSP grant RGP0060/2021, as well as ARC DP240100462, New Zealand Royal Society RDF 21-UOA-040, Marsden Grant 18-UOA-034, and University of Auckland, Faculty of Science Research Development Fund #3732317.

The authors acknowledge helpful discussions with colleagues including Jordan Douglas and Ashar Malik. The authors also wish to acknowledge the use of New Zealand eScience Infrastructure (NeSI) high performance computing facilities, consulting support and training services as part of this research. New Zealand’s national facilities are provided by NeSI and funded jointly by NeSI’s collaborator institutions and through the Ministry of Business, Innovation & Employment’s Research Infrastructure programme. URL https://www.nesi.org.nz.

## DATA AVAILABILITY STATEMENT

The data underlying this article and supplementary materials are available in the following GitHub repository: https://github.com/nmatzke/FullmerMatzke25/Fullmer_etal_2025

